# Comprehensive and quantitative analysis of white and brown adipose tissue by shotgun lipidomics

**DOI:** 10.1101/470690

**Authors:** Michal Grzybek, Alessandra Palladini, Vasileia I Alexaki, Michal A. Surma, Kai Simons, Triantafyllos Chavakis, Christian Klose, Ünal Coskun

## Abstract

Shotgun lipidomics enables an extensive analysis of lipids from tissues and fluids. Each specimen requires appropriate extraction and processing procedures to ensure good coverage and reproducible quantification of the lipidome. Adipose tissue (AT) has become a research focus with regard to its involvement in obesity-related pathologies. However, the quantification of the AT lipidome is particularly challenging due to the predominance of triacylglycerides, which elicit high ion suppression of the remaining lipid classes. We present a new and validated method for shotgun lipidomics of AT, which tailors the lipid extraction procedure to the target specimen and features high reproducibility with a linear dynamic range of at least 4 orders of magnitude for all lipid classes. Utilizing this method, we observed tissue-specific and diet-related differences in three AT types (brown, gonadal, inguinal subcutaneous) from lean and obese mice. Brown AT exhibited a distinct lipidomic profile with the greatest lipid class diversity and responded to high-fat diet by altering its lipid composition, which shifted towards that of white AT. Moreover, diet-induced obesity promoted an overall remodelling of the lipidome, where all three AT types featured a significant increase in longer and more unsaturated triacylglyceride and phospholipid species.

The here presented method facilitates reproducible systematic lipidomic profiling of AT and could be integrated with further –omics approaches used in (pre-)clinical research, in order to advance the understanding of the molecular metabolic dynamics involved in the pathogenesis of obesity-associated disorders.

## Introduction

Adipose tissue (AT) in mammals plays a critical role in systemic energy homeostasis[1-3]. Two main AT forms exist, the white adipose tissue (WAT), which functions primarily as an energy reservoir, and the brown adipose tissue (BAT), which contributes to thermoregulation, i.e. the production of heat to maintain body temperature stability during cold exposure through non-shivering thermogenesis. The expansion of AT observed in obesity is accompanied by local hypoxia[4-6], low-grade inflammation[7-11], development of systemic insulin resistance[5] and alterations in the secretion of fatty acids[12] or adipokines[4]. Thus, studying the regulation of lipid flux and metabolism in AT and changes thereof linked to obesity is important for understanding the central role of AT in obesity-related metabolic dysfunction.

Quantitative lipid analysis of biological samples by direct infusion mass spectrometry has proven to be a powerful tool for studying lipid metabolism[13-18]. It requires comparatively low sample amounts (e.g. 1 µl of plasma is sufficient to quantify a lipidome with comprehensive coverage[19]), relatively short measurement times - a prerequisite for the high-throughput analysis of large sample numbers, and exhibits excellent technical reproducibility. The use of internal standards enables direct quantification and application to a wide range of sample types, such as yeast[13 16], flies[20], *C. elegans[21]*, cellular samples[14 17 18] or various tissue and body fluids[19]. With regards to lipid quantification, ATs are particularly challenging because of their high content of neutral lipids (>99% triacylglycerols and cholesterol esters) that causes high ion suppression of the less abundant membrane lipids [9 22-24]. Previous approaches towards AT lipidomics analysis reported differences between BAT and WAT and demonstrated remodelling upon exercise or cold exposure. However, these studies applied generic methods of lipid extraction and measurement rather than techniques tailored to these tissues[23-26].

Here we present a shotgun lipidomics method for the analysis for ATs with an unprecedented coverage of more than 300 lipid species encompassing 20 lipid classes, and featuring high reproducibility with a linear dynamic range of at least 4 orders of magnitude for all lipid classes. We not only observe clear differences between BAT and WAT lipidomes, but also amongst WAT subtypes, i.e. gonadal (GAT) and inguinal subcutaneous (SAT) ATs. Furthermore, our protocol allows for assessing the impact of high-fat diet (HFD) on the remodelling of the lipidome, characterized by a significant increase in longer and more unsaturated acyl chains within the triacylglycerides and phospholipids.

## Results

### Experimental design, Method performance & standardization

Aiming at high reproducibility and standardization of the comprehensive analysis of AT lipids, we focused on developing a shotgun lipidomics method with an expanded dynamic range, high sensitivity, i.e. minimal limit of detection, and linearity. The starting material was AT (SAT, GAT and BAT) obtained from mice fed a control (CD) or high-fat diet (HFD). Typically, for preparation of tissue material for lipidomic analysis, samples are homogenised in aqueous buffers. However, following that standard procedure, we faced difficulties pipetting reproducible amounts of homogenised tissue material from the suspensions during aliquoting and dilution steps of the protocol, likely caused by the presence of large amount of fat droplets in homogenized AT samples. This reproducibility issue was solved by homogenization of AT in 50 vol% ethanol and subsequent dilution in pure ethanol (if required).

The reproducibility of the shotgun lipidomics method for AT was assessed by 6 repeated, independent measurements of identical aliquots of AT samples. For the assessment of reproducibility, only lipids present in at least 4 out of 6 replicates were considered. For the 276 lipids that fulfilled this requirement, the relative standard deviation (RSD) of each lipid molecule was calculated and plotted against its abundance (in pmol, **Figure 1B**) and the data revealed an inverse correlation between lipid abundance and RSD. The median technical variation is 6.8%, while 78.6% of the detected lipids show an RSD <15%. Taken together, the shotgun lipidomics approach provides a reproducible way to assess lipid abundance in the AT.

**Figure 1.**
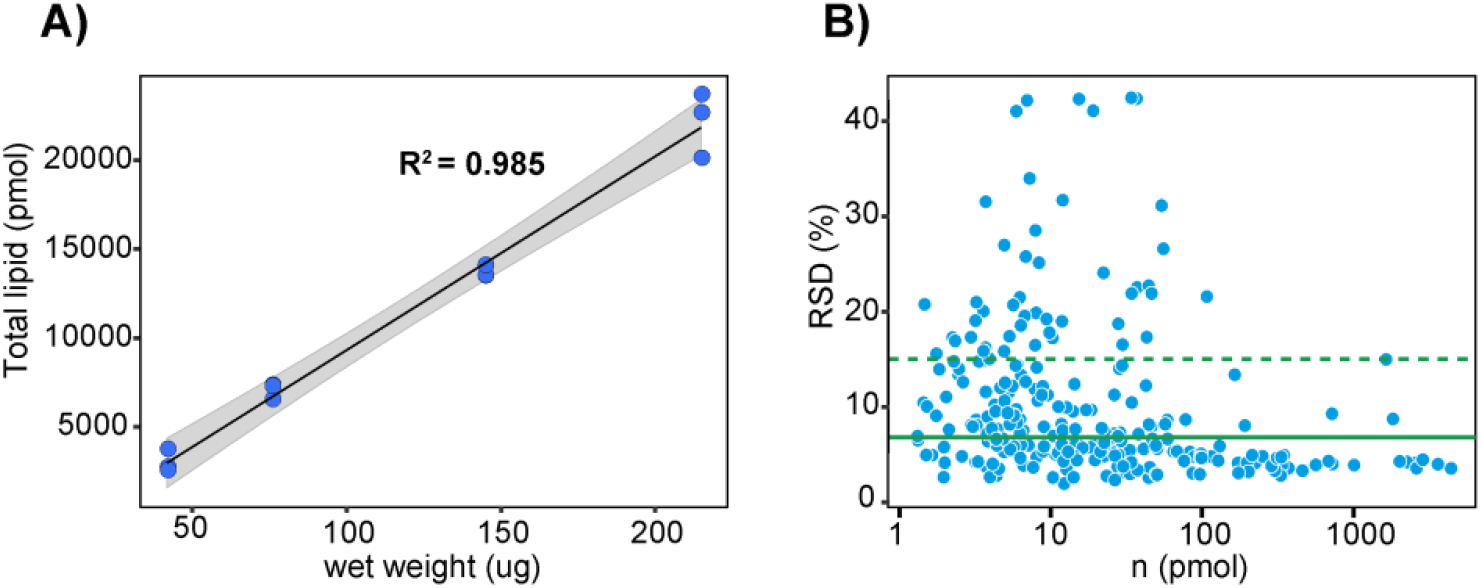
**A)** Linearity and proportionality assessed for total lipid content extracted from various amounts of SAT represented as linear regression of log-transformed lipid amounts and their intensities. **B)** Relative standard deviation (RSD) calculated for 276 lipids present in at least 4 out 6 replicates. Each lipid molecule RSD was calculated and plotted against its abundance (in pmol). The median technical variation is 6.8% (green solid line) and RSD <15% (green dotted line) was observed for 78.6% of the detected lipids.

To determine the optimal sample amount, a range of 35 to 200 µg of wet weight SAT were extracted in triplicate in the presence of fixed amounts of internal standards, and analysed by shotgun mass spectrometry. Total lipid content showed a linear response over the indicated range (**Figure 1**) and was proportional to sample amount, yielding an average of 98 pmol of lipid per µg wet weight. Furthermore, lipid species profiles became independent of sample amount above 70 µg of wet weight (**Table S1**).

The dynamic range, limit of detection (LOD) and limit of quantification (LOQ) are lipid class-specific parameters, which depend on differential extraction, ionisation and fragmentation efficiencies of the various lipid classes. In order to determine these parameters, lipid class-specific standards were titrated to a fixed amount of AT homogenate. Linearity and proportionality were assessed for 20 lipid classes by linear regression of log-transformed lipid amounts and their intensities, and reported as *R*^2^ and slope, respectively. Both, linearity and proportionality were excellent across a range of up to 4 orders of magnitude for all lipid classes, with values close to 1 for *R*^2^ and slope, respectively (**Table 1 & Figure S1**). LOD and LOQ were determined by weighted linear regression (with weights being 1/*χ*^2^) based on a signal-to-noise ratio of 3 for LOD and 10 for LOQ. For most lipid classes, LOD is around 1 pmol (Table 1). Provided that an optimal sample amount (100 µg of wet weight) of AT yields ∼10,000 pmol of total lipid (**Figure 1A**), 1 pmol of a given lipid molecule is theoretically detectable in a typical sample, which corresponds to 0.01 mol%.

**Table 1.** Limit of detection (LOD), Limit of quantification (LOQ), and linearity for various lipid classes in AT.

| class | min (pmol) | lod (pmol) | loq (pmol) | slope | R2 |
| --- | --- | --- | --- | --- | --- |
| CL | 0,23 | 0,83 | 2,52 | 0,96 | 0,99 |
| Cer | 0,09 | 0,67 | 2,04 | 0,94 | 1 |
| ST | 1,83 | 1,71 | 5,17 | 1,05 | 0,9 |
| HexCer | 0,09 | 0,93 | 2,81 | 0,94 | 0,99 |
| LPA | 0,09 | 0,81 | 2,46 | 1,04 | 0,99 |
| LPC | 0,23 | 0,70 | 2,11 | 0,97 | 0,99 |
| LPE | 0,09 | 0,78 | 2,38 | 0,94 | 0,99 |
| LPG | 0,09 | 0,92 | 2,80 | 0,94 | 0,99 |
| LPI | 0,09 | 1,03 | 3,13 | 1,01 | 0,98 |
| LPS | 0,09 | 1,42 | 4,32 | 0,99 | 0,92 |
| SM | 0,23 | 0,95 | 2,87 | 0,96 | 0,99 |
| TAG | 1,37 | 0,66 | 1,99 | 0,94 | 0,99 |
| CE | 1,37 | 3,24 | 9,83 | 0,88 | 0,94 |
| DAG | 0,09 | 0,90 | 2,72 | 0,89 | 0,99 |
| PA | 0,27 | 0,89 | 2,70 | 1,23 | 0,98 |
| PC | 0,91 | 0,92 | 2,79 | 0,96 | 0,99 |
| PE | 0,23 | 1,05 | 3,18 | 0,94 | 0,98 |
| PG | 0,09 | 0,79 | 2,38 | 1,03 | 0,99 |
| PI | 0,23 | 1,17 | 3,55 | 0,93 | 0,98 |
| PS | 0,23 | 1,1 | 3,32 | 1,11 | 0,97 |

### Tissue-specific and diet-related lipidomic trends

Using the newly validated shotgun lipidomics method, WAT (SAT and GAT) and BAT samples obtained from mice fed with either control or high-fat diet were assayed. As expected, the vast majority (∼99%) of lipids in all samples were triacylglycerides (TAG). Principal component analysis (PCA) shows that BAT segregates from SAT and GAT along the 1st principal component (PC1) (**Figure 2A**). This dimension, which explains about 28% of the variance, reveals that groups of samples differ mainly according to tissue type. Interestingly though, upon HFD the BAT lipidome shifts towards a more WAT-like composition, i.e. it approaches that of GAT and SAT samples from HFD-fed mice. In contrast, PC2 mainly explains diet-related differences. While both BAT clusters (CD and HFD) can be clearly distinguished, SAT and GAT samples overlap in the PCA reduced space. We therefore used a different explorative approach called Minimum Curvilinear Embedding (MCE)[27], an unsupervised machine learning for nonlinear dimensionality reduction (**Figure 2B**). Centered MCE was able to clearly segregate WAT samples on CD and, additionally, uncovered a trend of SAT and GAT samples to separate following HFD, though there was still a partial overlap. These findings are the consequence of a very limited variation between average SAT and GAT lipid species mol% in HFD cohorts, i.e. with a log2 fold change below 1 in almost all species (**Table S2**). In this experiment the core lipidome of AT, i.e. the species shared by all three tissue types on either diet, consisted of 204 lipids belonging to DAG (6), TAG (195), PC (1), PE (1), ST (1) (**Table S3**).

**Figure 2.**
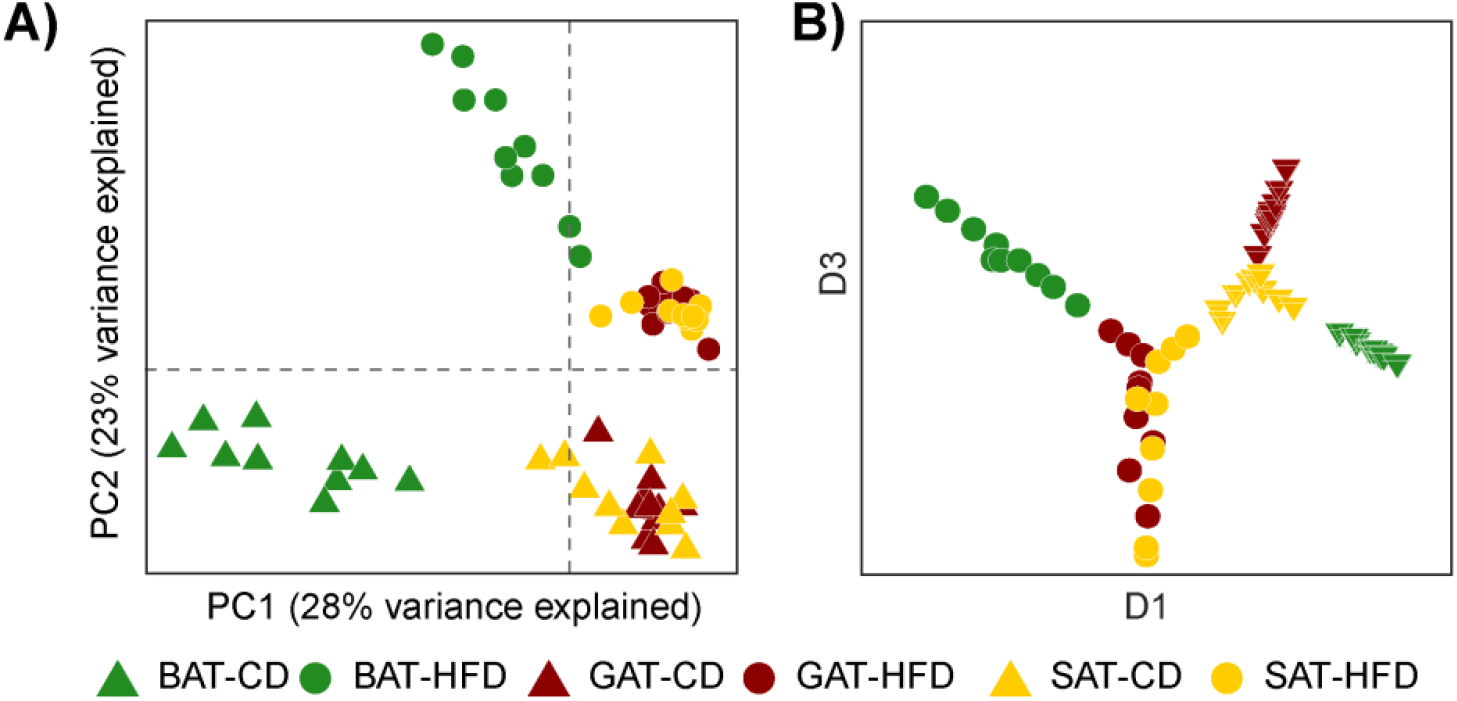
**A**) Principal Component Analysis (PCA) of AT lipidomes. BAT and WAT samples clearly and significantly separate, with GAT and SAT samples from CD-fed mice clustering separately from GAT and SAT samples from HFD-fed mice. PC1 significantly segregates samples according to tissue type, and PC2 significantly segregates samples according to diet. Adjusted p-values (Benjamini-Hochberg correction) for PC1 are: < 0.001 for BAT-CD vs. BAT-HFD; < 0.001 for BAT-CD and BAT-HFD vs. GAT and SAT on CD and HFD; > 0.05 for GAT-CD vs. GAT-HFD and vs. SAT-HFD, and for SAT-CD vs. SAT-HFD; nonsignificant for all other GAT vs. SAT comparisons. Comparisons along PC2 are significant except for GAT vs. SAT on the same diet. **B**) Centered Minimum Curvilinear Embedding (cMCE) on Spearman correlation. Dimensions 1 and 3 significantly differentiate almost all groups of samples. Adjusted p-values (Benjamini-Hochberg correction) for D1 are < 0.001 for all comparisons except for GAT-HFD vs. SAT-HFD where p-value < 0.01, and for GAT-CD vs. SAT-CD where p-value < 0.05. Along D3 all comparisons are significant except GAT-HFD vs. SAT-HFD.

PCA and MCE analyses showed that BAT lipidomes are particularly different from SAT and GAT; nevertheless, in all three fat depots TAGs were the major lipid class present, reaching 98-99% of all identified lipids regardless of diet. By excluding TAG lipids from the analysis, we focused on changes concerning the less abundant classes of lipids. Among these, DAGs were the most abundant in WAT (**Figure 3**), a finding likely associated with lipid metabolism and TAG turnover. In BAT, DAG levels are similar to those of PC and PE and other lipid classes like CL, LPC, PI and PG that were detected in BAT but remained undetectable in WAT. Importantly, cardiolipins (CL) are present in BAT on both diets, while absent in both SAT and GAT (**Figure 3 and Table S4**), most likely linked to the larger number of mitochondria in BAT than in WAT[28]. On CD, BAT contains higher amounts of glycerophospholipids and less cholesterol than WAT. On HFD though, cholesterol tends to increase in BAT while it shows a significant decrease in SAT and GAT.

**Figure 3.**
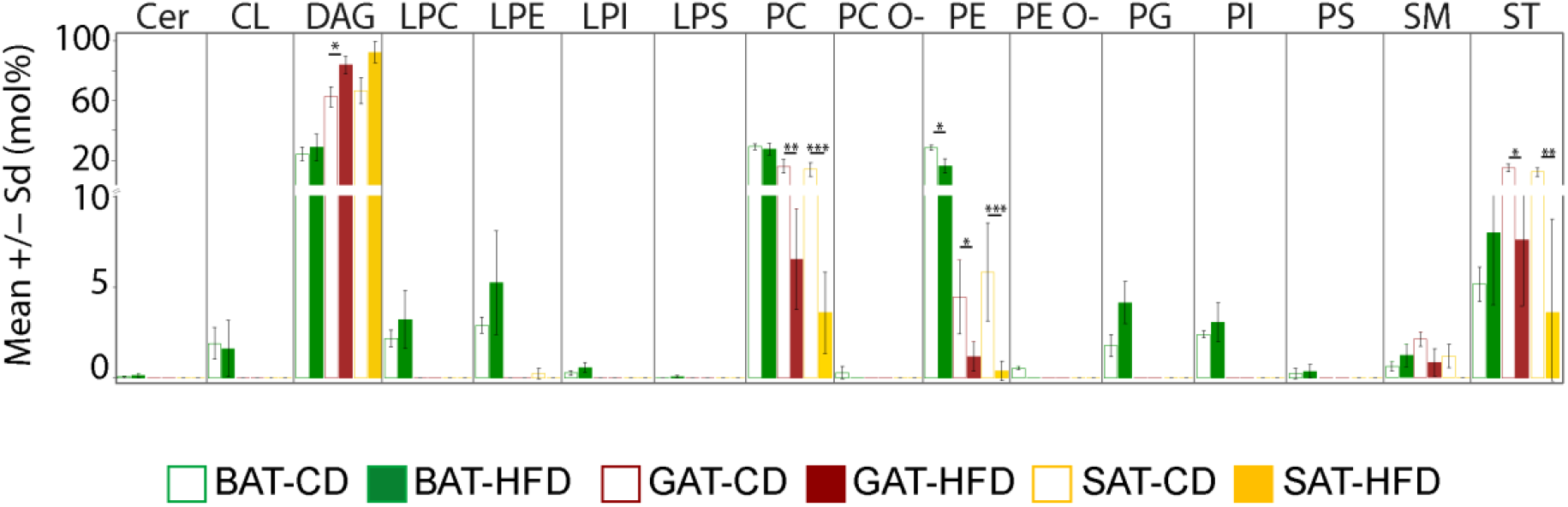
Lipid class composition of white and brown adipose tissues following chow and high-fat diet excluding TAGs. Asterisks indicate a significant difference between CD and HFD (p-value < 0.05*, < 0.01**, < 0.001***). The different depot types are color coded as follows: BAT green, GAT dark red, SAT yellow. White bars with colored border indicate cohorts on CD, colored bars indicate cohorts on HFD.

Following HFD, the acyl chain composition of TAGs (**Figure 4A & S2A**) in all three AT depots shows a significant decrease of species containing a cumulative 48-50 carbon atoms in their acyl chains, while TAGs with 54-56 carbon atoms increase. Interestingly in BAT as well, TAGs containing 52 carbon atoms decrease significantly, while no substantial changes are observed in SAT or GAT. The increase in length correlates with the increase in the number of double bonds: TAGs containing a total of 0, 1 or 2 double bonds in their acyl chains are more abundant following CD than HFD, where most TAGs contain a total number of 4 and more double bonds. TAGs with 3 double bonds display different trends in brown and white AT. In BAT they significantly increase upon HFD, reaching the average amount displayed by white depots, but they do not change in GAT, whereas in SAT they decrease significantly. In summary, TAGs become longer and more unsaturated when animals are on HFD and these changes are more pronounced in BAT than in WAT.

**Figure 4.**
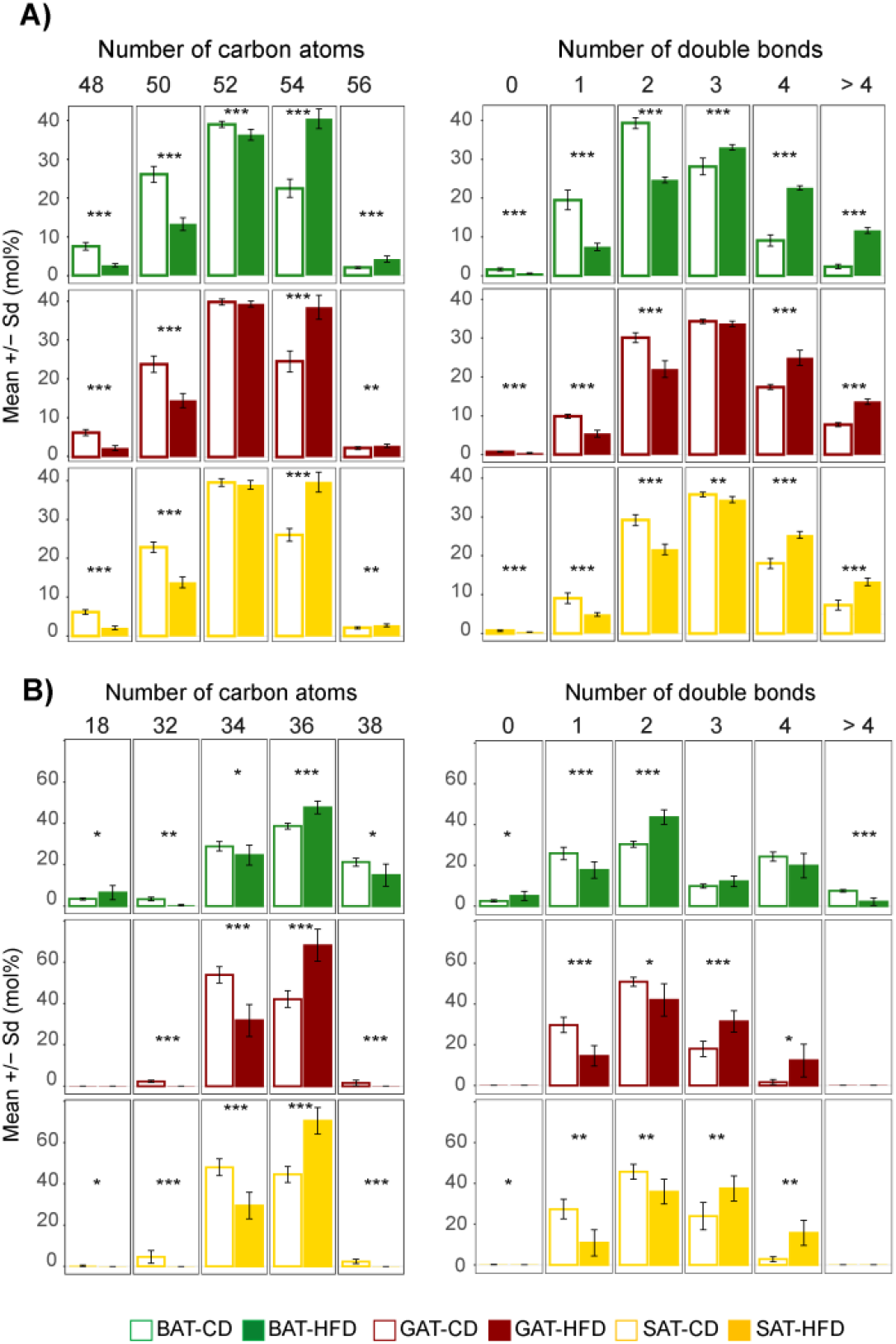
Total acyl chain length (left side) and unsaturation (right side) profiles of: A) TAGs; B) membrane lipids. Lipids were regrouped according to the number of carbon atoms and the number of double bonds present in their acyl chains. Mean and standard deviation were calculated for each cohort using mol% - transformed data. Only length groups > 3 mol% are represented; complete acyl chain length profiles of A and B are shown in the supplementary Figure S3. Asterisks indicate a significant difference between CD and HFD (p-value < 0.05*, < 0.01**, < 0.001***). The different types of AT are color-coded as follows: BAT green, GAT dark red, SAT yellow. White bars with coloured border indicate cohorts on CD; coloured bars indicate cohorts on HFD.

Remarkably, the changes observed for TAGs are hardly reflected in the acyl chain composition of membrane lipids. Upon HFD, lipids containing 36 carbon atoms significantly increase in abundance, while the abundance of other lipids, both shorter and longer than 36, decreases regardless of the AT depot type (**Figure 4B & S2B**). The only exception concerns lipids containing 18 carbon atoms: these are lysolipids, and they slightly increase in BAT. The double bond composition displays different trends for white and brown AT, except for monounsaturated lipids, which always decrease on HFD. In white AT, saturated and di-unsaturated lipids decrease upon HFD, while in BAT they increase, and lipids containing 3 or 4 double bonds increase, while in BAT they do not change significantly. Polyunsaturated lipids (> 4 double bonds) are detected only in BAT, where they significantly decrease on HFD. In WAT depots, the length and unsaturation profiles of membrane lipids and triglycerides follow the same basic rules, as the relative increase in acyl chain length is accompanied by an increase in the number of double bonds. In contrast, in BAT membrane lipids the unsaturation pattern differs from that of triglycerides, in that we observe an inverse behavior of mono- and di-unsaturated species on HFD, and a concomitant clear decrease of highly poly-unsaturated species (mean values and standard deviation of length and unsaturation groups are given in **Table S5**).

To further understand the diet-dependent changes affecting the lipidome of the various AT depots, we inspected the difference between the means (HFD-CD) of the single subspecies in membrane and storage lipidomes (**Figure 5** and **S3**). By defining a threshold difference of |0.5| mol%, we excluded lipids with a mean mol% < 0.5 in both diets. This highlights the largest differences concerning highly abundant features (i. e. with a mean value on either diet near or above the 75^th^ percentile). The levels of DAGs containing two C18-long acyl chains, e.g. octadecenoic (18:1), octadecadienoic (18:2) and octadecatrienoic (18:3) FA, increase upon HFD in both WAT and BAT, and, simultaneously, DAGs containing either hexadecanoic (16:0) or hexadecenoic (16:1) FA decrease (**Figure 5 A-C**). A clear exception is DAG 16:0-18:2, which is more abundant in all three AT depots on HFD. It should be noted that in GAT and SAT, lipids with a total length of 36 carbons exclusively belong to the DAGs, whereas in BAT samples, C36 species also occur in other lipid classes (PG, PI, PC). A major difference in the behaviour of BAT and WAT during HFD is the abundance of cholesterol. In WAT cholesterol is the major lipid and decreases upon HFD, whereas it accumulates in BAT upon HFD. Interestingly, the lipids that display the highest decrease upon HFD in BAT are three PE species containing eicosatetraenoic acid (16:0/20:4; 18:0/20:4; 18:1/20:4).

**Figure 5.**
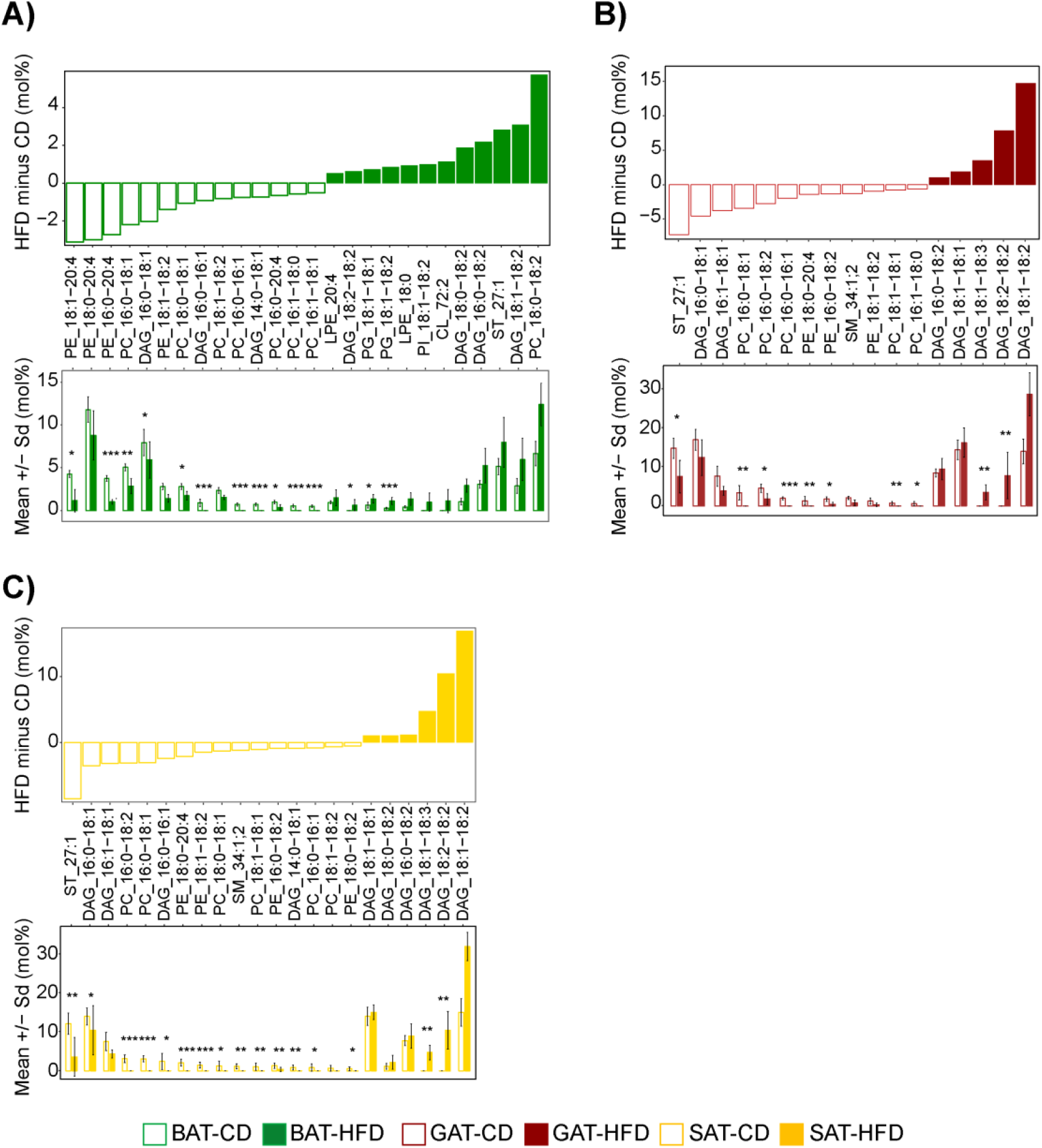
Subspecies level analysis of membrane lipids in fat tissues Upper panel bar plots represent the difference between the means of HFD and CD in the respective adipose tissues: A) BAT; B) GAT; C) SAT. Only species with ∆ > |0.5| mol% are shown. Lower panel bar plots represent the mean amount and standard deviation of the subspecies in CD and HFD cohorts. Asterisks indicate the species that differ significantly between diets according to the pairwise Mann-Whitney test performed on the log-transformed dataset; p-values were adjusted according to the Benjamini-Hochberg correction for multiple comparisons.

The results for TAGs (**Figure S3 A-C**) support the length and unsaturation profiles shown in **Figure 4**. Specifically, the greatest differences concern shorter (C50 and C52 total length), less unsaturated subspecies in samples on CD, and longer (C52 and C54), more unsaturated subspecies in samples on HFD. Additional information is given by the acyl chain identified within each TAG: the increased abundance of C54 species in samples on HFD mostly concerns TAGs where the identified acyl chain comprises 18C.

The differences in lipid composition between BAT and WAT are easily distinguished by both PCA and MCE. However, differences between WAT depots are much more difficult to discern, and only MCE succeeded in distinguishing GAT and SAT on both diets, remarkably identifying patterns guided by ‘private’ features, i.e. lipids exclusive to a given cohort (Table S6). Nonetheless, when focusing on the core lipidome, even MCE cannot segregate GAT and SAT, since abundance patterns are too similar on both diets (**Figure S3, Table S3**).

## Discussion

Shotgun lipidomics allows for parallel profiling of hundreds of structurally and functionally diverse lipids. Nevertheless, obtaining a comprehensive, reproducible and quantitative lipidomics analysis of AT is particularly challenging. Here, we thoroughly characterized a shotgun lipidomics method for AT analysis and its validation with respect to sample amounts, linearity, sensitivity and reproducibility (**Figure 1 & Table 1**). This method requires about 100 µg of AT to yield about 10,000 pmol total lipid, which represents an optimal amount for shotgun lipidomics. It allows the detection of more than 300 individual lipid molecules, belonging to 20 different lipid classes, with a high dynamic range of about 4 orders of magnitude and sensitivity in the sub-µM range, and high reproducibility below 10% RSD. By quantitatively inspecting significant differences between the lipidomes of WAT and BAT from mice fed with control and high-fat diets, we provide proof of concept and validation. At the present stage, we cannot provide a mechanistic explanation of the biological processes underlying these changes, likely to be associated with the nature of the fatty acids consumed in the diet. We hypothesise that they are coupled to the physiological mechanisms by which adipocyte membranes respond to AT expansion, associated to positive energy balance ultimately leading to obesity.

In HFD-fed mice, we observed an accumulation of TAGs with a total length of 54 carbons and a concomitant decrease of TAGs with 48-50 carbons (**Figure 4A**), meaning that 16-carbon acyl chains are replaced by 18-carbon acyl chains (**Figure S2**). Similarly, we observed an overall accumulation of non-storage lipids made of acyl chains containing 18 carbons and including various numbers of double bonds. In this regard, we noticed a relative increase of DAG, PC and PE species with 18:0, 18:1 and 18:2 acyl chains, alongside a decrease of species containing the 16-carbon acyl chain (**Figure 5**). These observations are in agreement with a previously published study on twins discordant for obesity, which suggested that obesity-related changes involve accumulation of membrane lipids containing longer and more unsaturated fatty acids than in the lean individuals[22]. Such a fatty-acyl composition was suggested to sustain optimal membrane physicochemical properties during adipocyte expansion.

In comparison to other tissue types the primary function of AT is to accumulate storage lipids is. Excess lipids, however, are stored ectopically in other tissues such as muscle, liver, and pancreas, a situation culminating to metabolic dysfunction. Indeed, the expansion of AT in obesity has effects that go beyond body-weight gain, usually associated with increased levels of circulating FAs, and inflammatory mediators that are now recognized as a major cause of decreased insulin sensitivity. Interestingly, many cohort studies reported that some obese individuals display a phenotype referred to as ‘‘metabolically healthy obesity’’, recognized by insulin sensitivity, smaller fat cells and less pronounced AT inflammation than those observed in insulin-resistant obese individuals[29-31]. It is also established that the two obesity phenotypes differ in SAT gene expression, AMPK activity or oxidative stress[32-34]. Thus, profiling AT lipidomes can advance knowledge of the processes underlying lipid metabolism and resolve differences of AT hypertrophy that correlate with the maintenance of a metabolically healthy status. The protocol presented here precisely provides the platform for addressing this issue.

Some fat depots are implicated more than others in the development of insulin resistance[35 36]. Increasing the number of samples and identifying all three acyl chains of TAGs would shed light on the differences characterizing GAT and SAT, and correlate diet-related lipidomic patterns to metabolic parameters. This would pave the way to a fundamental understanding of the dynamics underlying hypertrophy of distinct WAT depots and the development of metabolic syndrome.

Lipidomics analysis can also supply additional information to studies on the physiology of BAT and the beiging of WAT. In fact, our method could measure the lipidomes of both AT depots from lean and obese individuals, resolving the differences among the tested groups. Besides generating the distinct profiles of WAT and BAT lipidomes, as previously shown by others[23], the data obtained in this study also provide a snapshot of the diverse response of brown and white AT to overnutrition (**Figures 3 & 4**). As shown by PCA (Figure 3A), BAT on HFD shifts toward a more WAT-like composition, despite maintaining its identity, since it still clusters separately from WAT samples. BAT’s distinct lipidomic profile under HFD is achieved already at class level: its number of lipid classes, higher than that of WAT, remains unchanged on HFD (**Figure 3**). Additionally, we do not observe any clear change in the total levels of DAGs, a completely different behaviour from that exhibited by WAT depots, where DAG levels significantly increase. On the other hand, in BAT on HFD, PE levels significantly drop, while cholesterol, PG, PI and SM increase. The similarities of BAT and WAT lipidomes following HFD are visible in the unsaturation and length profiles, where amounts of lipids with the same length or number of double bonds converge, independent of their initial differences on CD (**Figure 4 & S2**). The important and beneficial role of BAT in fatty acid oxidation and thermogenesis has mainly been shown in animal models[26], but recent evidence from human BAT studies was also published[2]. Furthermore, in specific cases human SAT undergoes browning/beiging, associated with an increased whole-body metabolic rate[37]. The interest in increasing BAT metabolism or promoting browning/beiging of WAT in humans as a potential strategy for the treatment of obesity and its related metabolic disorder is enormous. However, investigating the physiological differences of the various AT depots requires a systematic lipidomic analysis as an essential element of the -omics pipeline. This would provide core data towards elucidating AT remodelling mechanisms and thus potentially facilitate the development of therapeutic approaches against obesity-induced metabolic diseases.

The presented method can easily be utilized for non-AT samples containing high amounts of neutral lipids, e.g. tissue samples from subjects with fatty-liver disease [38 39]. Such samples, under common extraction and measurement conditions, would interfere with full lipidomic analysis due to ion suppression. Obtaining detailed lipidomic profiles from patients at various stages of a disease will be crucial for investigating the aetiology of obesity-related pathologies. Samples rich in lipid droplets are another example[40-42]. Studying the biogenesis and metabolism of these organelles or lipid droplet-associated organelles like ER[43] or mitochondria[44] would benefit from monitoring the dynamics of full lipid profiles within the isolated samples. The presented approach offers a solution to handle these challenges.

## Materials & Methods

### Chemicals

Water, propan-2-ol, and methanol were purchased from Fischer Scientific. Methyl tert-butyl ether, chloroform, ammonium bicarbonate and ammonium acetate were purchased from Sigma–Aldrich. All chemicals were analytical grade. Synthetic lipid standards were purchased from Avanti Polar Lipids, Larodan Fine Chemicals, and Sigma–Aldrich.

### Feeding of mice

Six-weeks-old female C57BL/6 mice were fed a control diet (CD, D12450B, Research Diets, NJ, USA, 20% protein, 70% carbohydrates, 10% fat), or a high fat diet (HFD, D12492, Research Diets, NJ, USA, 20% protein, 20% carbohydrates, 60% fat) for 20 weeks, as previously described[6 10 45]. Experiments were approved by the Landesdirektion Sachsen, Germany.

### Sample preparation

ATs were homogenized directly in a Turrax homogenizer for 60 s in 0.5 ml of ice-cold 150 mM ammonium bicarbonate and ethanol (50/50 v/v). Homogenates were subsequently diluted 1:10 (v/v) in ethanol and used for lipid extraction.

### Lipid extraction

Mass spectrometry-based lipid analysis was performed at Lipotype GmbH (Dresden, Germany) as described[14]. Lipids were extracted using a two-step chloroform/methanol procedure[13]. Samples were spiked with internal lipid standard mixture containing: 30 pmol cardiolipin 16:1/15:0/15:0/15:0 (CL), 30 pmol ceramide 18:1;2/17:0 (Cer), 100 pmol diacylglycerol 17:0/17:0 (DAG), 30 pmol hexosylceramide 18:1;2/12:0 (HexCer), 30 pmol lyso-phosphatidate 17:0 (LPA), 50 pmol lyso-phosphatidylcholine 12:0 (LPC), 30 pmol lyso-phosphatidylethanolamine 17:1 (LPE), 30 pmol lyso-phosphatidylglycerol 17:1 (LPG), 20 pmol lyso-phosphatidylinositol 17:1 (LPI), 30 pmol lyso-phosphatidylserine 17:1 (LPS), 50 pmol phosphatidate 17:0/17:0 (PA), 150 pmol phosphatidylcholine 17:0/17:0 (PC), 75 pmol phosphatidylethanolamine 17:0/17:0 (PE), 50 pmol phosphatidylglycerol 17:0/17:0 (PG), 50 pmol phosphatidylinositol 16:0/16:0 (PI), 100 pmol phosphatidylserine 17:0/17:0 (PS), 100 pmol cholesterol ester 20:0 (CE), 50 pmol sphingomyelin 18:1;2/12:0;0 (SM), 500 pmol triacylglycerol 17:0/17:0/17:0 (TAG) and 300 pmol cholesterol D6 (Chol). After extraction, the organic phase was transferred to an infusion plate and dried in a speed vacuum concentrator. 1st step dry extract was re-suspended in 7.5 mM ammonium acetate in chloroform/methanol/propanol (1:2:4; V:V:V) and 2nd step dry extract in 33% ethanol solution of methylamine in chloroform/methanol (0.003:5:1; V:V:V). All liquid handling steps were performed using Hamilton Robotics STARlet robotic platform with the Anti Droplet Control feature for organic solvents pipetting.

### Mass spectrometry

Samples were analysed by direct infusion on a QExactive mass spectrometer (Thermo Scientific) equipped with a TriVersa NanoMate ion source (Advion Biosciences). Samples were analysed in both positive and negative ion modes with a resolution of R_m/z=200_=280000 for MS and R_m/z=200_=17500 for MSMS experiments, in a single acquisition. MSMS was triggered by an inclusion list encompassing corresponding MS mass ranges scanned in 1 Da increments[19]. Both MS and MSMS data were combined to monitor CE, DAG and TAG ions as ammonium adducts; PC, PC O-, as acetate adducts; and CL, PA, PE, PE O-, PG, PI and PS as deprotonated anions. MS only was used to monitor LPA, LPE, LPE O-, LPI and LPS as deprotonated anions; Cer, HexCer, SM, LPC and LPC O- as acetate adduct and cholesterol as ammonium adduct of an acetylated derivative[46].

### Data analysis and post-processing

Data were analysed with in-house developed lipid identification software based on LipidXplorer[47 48]. Data post-processing and normalization were performed using an in-house developed data management system. If not stated otherwise, only lipid identification with a signal-to-noise ratio > 5, a signal intensity 5-fold higher than in corresponding blank samples, and lipids present in at least 5 out of 10 replicates was considered for further data analysis.

Data were analysed with R version 3.5 (R Core Team, 2018) using tidyverse packages version 1.1.1[49] and plots were created with ggplot2 version 2.2.1[50].

### Lipid nomenclature

When describing different lipid species the following annotations are used: Lipid class-[sum of carbon atoms]:[sum of double bonds];[sum of hydroxyl groups], i.e. SM-34:1;2 means an SM species with 34 carbon atoms, 1 double bond and 2 hydroxyl groups in the ceramide backbone. Lipid subspecies annotation contains additional information on the exact identity of their fatty acids. For example, PI-34:1;0(18:1;0-16:0;0) denotes phosphatidylinositol with a total length of its fatty acids equal to 34 carbon atoms, total number of double bonds in its fatty acids equal to 1 and 0 hydroxylations with C18:1 (oleic) and C16:0 (palmitic) fatty acids. When the exact position of fatty acids in relation to the glycerol backbone (*sn1* or *sn2*) is not discernible an underscore “_” separating the acyl chains is used. In case the *sn* position is known, acyl chain information is separated by a slash “/”. TAG features are reported as pairs of intact ion and neutral loss of a fatty acid. For example, TAG-50:1;0-FA-18:1;0 refers to the neutral loss of an 18:1 fatty acid belonging to the intact TAG-50:1;0 molecule.

### Descriptive statistics of lipidomics data

Lipid features present in less than 50% of the samples in a cohort were filtered in that given cohort. Analyses were performed in R (R Core Team 2018) on the mol%-transformed dataset, i.e., after transforming raw data (picomol) to mole percent (each quantity was divided by the sum of the lipids detected in its respective sample and multiplied by 100). Data structure was analysed by means of Principal Component Analysis (PCA) using the Singular Value Decomposition function, and by means of the nonlinear machine learning called Minimum Curvilinear Embedding (MCE)[27]. Total carbon chain length and unsaturation plots result from grouping together all the lipids that present the same number of carbon atoms (total length) or the same number of double bonds (unsaturation) and calculating their mean and standard deviation in each cohort of samples. The difference between the means was calculated for each species by subtracting the mean of the controls from the mean of the treated samples (Treated *minus* Reference). Significance was calculated by means of the non-parametric Wilcoxon test and p-values were adjusted after the Benjamini-Hochberg correction. Alongside R-base functions the following packages were used: reshape2[49] and ggplot2[50].

## Acknowledgments

We thank Cornelia Schroeder and Sider Penkov for critical reading of the manuscript. CK and MAS would like to thank Steffi Lenhard for excellent technical assistance. TC was supported by the ERC (DEMETINL).

## References

1. Lempradl A, Pospisilik JA, Penninger JM. Exploring the emerging complexity in transcriptional regulation of energy homeostasis. Nat Rev Genet 2015;16(11):665-81 doi:10.1038/nrg3941[published Online First: Epub Date]|.

2. Chondronikola M, Volpi E, Borsheim E, et al. Brown adipose tissue improves whole-body glucose homeostasis and insulin sensitivity in humans. Diabetes 2014;63(12):4089-99 doi:10.2337/db14-0746[published Online First: Epub Date]|.

3. Choe SS, Huh JY, Hwang IJ, Kim JI, Kim JB. Adipose Tissue Remodeling: Its Role in Energy Metabolism and Metabolic Disorders. Front Endocrinol (Lausanne) 2016;7:30 doi:10.3389/fendo.2016.00030[published Online First: Epub Date]|.

4. Engin A. Adipose Tissue Hypoxia in Obesity and Its Impact on Preadipocytes and Macrophages: Hypoxia Hypothesis. Adv Exp Med Biol 2017;960:305-26 doi:10.1007/978-3-319-48382-5_13[published Online First: Epub Date]|.

5. Hosogai N, Fukuhara A, Oshima K, et al. Adipose tissue hypoxia in obesity and its impact on adipocytokine dysregulation. Diabetes 2007;56(4):901-11 doi:10.2337/db06-0911[published Online First: Epub Date]|.

6. Garcia-Martin R, Alexaki VI, Qin N, et al. Adipocyte-Specific Hypoxia-Inducible Factor 2alpha Deficiency Exacerbates Obesity-Induced Brown Adipose Tissue Dysfunction and Metabolic Dysregulation. Mol Cell Biol 2016;36(3):376-93 doi:10.1128/MCB.00430-15[published Online First: Epub Date]|.

7. Reilly SM, Saltiel AR. Adapting to obesity with adipose tissue inflammation. Nat Rev Endocrinol 2017 doi:10.1038/nrendo.2017.90[published Online First: Epub Date]|.

8. Hong CP, Yun CH, Lee GW, Park A, Kim YM, Jang MH. TLR9 regulates adipose tissue inflammation and obesity-related metabolic disorders. Obesity (Silver Spring) 2015;23(11):2199-206 doi:10.1002/oby.21215[published Online First: Epub Date]|.

9. Sedger LM, Tull DL, McConville MJ, et al. Lipidomic Profiling of Adipose Tissue Reveals an Inflammatory Signature in Cancer-Related and Primary Lymphedema. PLoS One 2016;11(5):e0154650 doi:10.1371/journal.pone.0154650[published Online First: Epub Date]|.

10. Chung KJ, Chatzigeorgiou A, Economopoulou M, et al. A self-sustained loop of inflammation-driven inhibition of beige adipogenesis in obesity. Nat Immunol 2017;18(6):654-64 doi:10.1038/ni.3728[published Online First: Epub Date]|.

11. Chung KJ, Nati M, Chavakis T, Chatzigeorgiou A. Innate immune cells in the adipose tissue. Rev Endocr Metab Disord 2018 doi:10.1007/s11154-018-9451-6[published Online First: Epub Date]|.

12. Mittendorfer B. Origins of metabolic complications in obesity: adipose tissue and free fatty acid trafficking. Curr Opin Clin Nutr Metab Care 2011;14(6):535-41 doi:10.1097/MCO.0b013e32834ad8b6[published Online First: Epub Date]|.

13. Ejsing CS, Sampaio JL, Surendranath V, et al. Global analysis of the yeast lipidome by quantitative shotgun mass spectrometry. 2009;106(7):2136

14. Sampaio JL, Gerl MJ, Klose C, et al. Membrane lipidome of an epithelial cell line. Proceedings of the National Academy of Sciences of the United States of America 2011;108(5):1903-07 doi:10.1073/pnas.1019267108[published Online First: Epub Date]|.

15. Kjellqvist S, Klose C, Surma MA, et al. Identification of Shared and Unique Serum Lipid Profiles in Diabetes Mellitus and Myocardial Infarction. J Am Heart Assoc 2016;5(12) doi:10.1161/JAHA.116.004503[published Online First: Epub Date]|.

16. Surma MA, Klose C, Peng D, et al. A lipid E-MAP identifies Ubx2 as a critical regulator of lipid saturation and lipid bilayer stress. Mol Cell 2013;51(4):519-30 doi:10.1016/j.molcel.2013.06.014[published Online First: Epub Date]|.

17. Mitroulis I, Ruppova K, Wang B, et al. Modulation of Myelopoiesis Progenitors Is an Integral Component of Trained Immunity. Cell 2018;172(1-2):147-61 e12 doi:10.1016/j.cell.2017.11.034[published Online First: Epub Date]|.

18. Stefanko A, Thiede C, Ehninger G, Simons K, Grzybek M. Lipidomic approach for stratification of acute myeloid leukemia patients. PLoS One 2017;12(2):e0168781 doi:10.1371/journal.pone.0168781[published Online First: Epub Date]|.

19. Surma MA, Herzog R, Vasilj A, et al. An automated shotgun lipidomics platform for high throughput, comprehensive, and quantitative analysis of blood plasma intact lipids. Eur J Lipid Sci Technol 2015;117(10):1540-49 doi:10.1002/ejlt.201500145[published Online First: Epub Date]|.

20. Carvalho M, Sampaio JL, Palm W, Brankatschk M, Eaton S, Shevchenko A. Effects of diet and development on the Drosophila lipidome. Mol Syst Biol 2012;8:600 doi:10.1038/msb.2012.29[published Online First: Epub Date]|.

21. Prasain JK, Wilson L, Hoang HD, Moore R, Miller MA. Comparative Lipidomics of Caenorhabditis elegans Metabolic Disease Models by SWATH Non-Targeted Tandem Mass Spectrometry. Metabolites 2015;5(4):677-96 doi:10.3390/metabo5040677[published Online First: Epub Date]|.

22. Pietilainen KH, Rog T, Seppanen-Laakso T, et al. Association of lipidome remodeling in the adipocyte membrane with acquired obesity in humans. PLoS Biol 2011;9(6):e1000623 doi:10.1371/journal.pbio.1000623[published Online First: Epub Date]|.

23. May FJ, Baer LA, Lehnig AC, et al. Lipidomic Adaptations in White and Brown Adipose Tissue in Response to Exercise Demonstrate Molecular Species-Specific Remodeling. Cell Rep 2017;18(6):1558-72 doi:10.1016/j.celrep.2017.01.038[published Online First: Epub Date]|.

24. Baker RC, Nikitina Y, Subauste AR. Analysis of Adipose Tissue Lipid Using Mass Spectrometry. Method Enzymol 2014;538:89-105 doi:10.1016/B978-0-12-800280-3.00006-2[published Online First: Epub Date]|.

25. Caesar R, Manieri M, Kelder T, et al. A combined transcriptomics and lipidomics analysis of subcutaneous, epididymal and mesenteric adipose tissue reveals marked functional differences. PLoS One 2010;5(7):e11525 doi:10.1371/journal.pone.0011525[published Online First: Epub Date]|.

26. Marcher AB, Loft A, Nielsen R, et al. RNA-Seq and Mass-Spectrometry-Based Lipidomics Reveal Extensive Changes of Glycerolipid Pathways in Brown Adipose Tissue in Response to Cold. Cell Rep 2015;13(9):2000-13 doi:10.1016/j.celrep.2015.10.069[published Online First: Epub Date]|.

27. Cannistraci CV, Ravasi T, Montevecchi FM, Ideker T, Alessio M. Nonlinear dimension reduction and clustering by Minimum Curvilinearity unfold neuropathic pain and tissue embryological classes. Bioinformatics 2010;26(18):i531-9 doi:10.1093/bioinformatics/btq376[published Online First: Epub Date]|.

28. Ito T, Tanuma Y, Yamada M, Yamamoto M. Morphological studies on brown adipose tissue in the bat and in humans of various ages. Arch Histol Cytol 1991;54(1):1-39

29. Primeau V, Coderre L, Karelis AD, et al. Characterizing the profile of obese patients who are metabolically healthy. Int J Obes (Lond) 2011;35(7):971-81 doi:10.1038/ijo.2010.216[published Online First: Epub Date]|.

30. Sims EA. Are there persons who are obese, but metabolically healthy? Metabolism 2001;50(12):1499-504 doi:10.1053/meta.2001.27213[published Online First: Epub Date]|.

31. Samocha-Bonet D, Chisholm DJ, Tonks K, Campbell LV, Greenfield JR. Insulin-sensitive obesity in humans - a ‘favorable fat’ phenotype? Trends Endocrinol Metab 2012;23(3):116-24 doi:10.1016/j.tem.2011.12.005[published Online First: Epub Date]|.

32. Xu XJ, Pories WJ, Dohm LG, Ruderman NB. What distinguishes adipose tissue of severely obese humans who are insulin sensitive and resistant? Curr Opin Lipidol 2013;24(1):49-56 doi:10.1097/MOL.0b013e32835b465b[published Online First: Epub Date]|.

33. Elbein SC, Kern PA, Rasouli N, Yao-Borengasser A, Sharma NK, Das SK. Global gene expression profiles of subcutaneous adipose and muscle from glucose-tolerant, insulin-sensitive, and insulin-resistant individuals matched for BMI. Diabetes 2011;60(3):1019-29 doi:10.2337/db10-1270[published Online First: Epub Date]|.

34. Qatanani M, Tan Y, Dobrin R, et al. Inverse regulation of inflammation and mitochondrial function in adipose tissue defines extreme insulin sensitivity in morbidly obese patients. Diabetes 2013;62(3):855-63 doi:10.2337/db12-0399[published Online First: Epub Date]|.

35. Sethi JK, Vidal-Puig AJ. Thematic review series: adipocyte biology. Adipose tissue function and plasticity orchestrate nutritional adaptation. J Lipid Res 2007;48(6):1253-62 doi:10.1194/jlr.R700005-JLR200[published Online First: Epub Date]|.

36. Wajchenberg BL, Giannella-Neto D, da Silva ME, Santos RF. Depot-specific hormonal characteristics of subcutaneous and visceral adipose tissue and their relation to the metabolic syndrome. Horm Metab Res 2002;34(11-12):616-21 doi:10.1055/s-2002-38256[published Online First: Epub Date]|.

37. Sidossis LS, Porter C, Saraf MK, et al. Browning of Subcutaneous White Adipose Tissue in Humans after Severe Adrenergic Stress. Cell Metab 2015;22(2):219-27 doi:10.1016/j.cmet.2015.06.022[published Online First: Epub Date]|.

38. Feng S, Gan L, Yang CS, et al. Effects of Stigmasterol and beta-Sitosterol on Nonalcoholic Fatty Liver Disease in a Mouse Model: A Lipidomic Analysis. J Agric Food Chem 2018;66(13):3417-25 doi:10.1021/acs.jafc.7b06146[published Online First: Epub Date]|.

39. Puri P, Baillie RA, Wiest MM, et al. A lipidomic analysis of nonalcoholic fatty liver disease. Hepatology 2007;46(4):1081-90 doi:10.1002/hep.21763[published Online First: Epub Date]|.

40. Hartler J, Kofeler HC, Trotzmuller M, Thallinger GG, Spener F. Assessment of lipidomic species in hepatocyte lipid droplets from stressed mouse models. Sci Data 2014;1:140051 doi:10.1038/sdata.2014.51[published Online First: Epub Date]|.

41. Chitraju C, Trotzmuller M, Hartler J, et al. Lipidomic analysis of lipid droplets from murine hepatocytes reveals distinct signatures for nutritional stress. J Lipid Res 2012;53(10):2141-52 doi:10.1194/jlr.M028902[published Online First: Epub Date]|.

42. Zhi Y, Taylor MC, Campbell PM, et al. Comparative Lipidomics and Proteomics of Lipid Droplets in the Mesocarp and Seed Tissues of Chinese Tallow (Triadica sebifera). Front Plant Sci 2017;8:1339 doi:10.3389/fpls.2017.01339[published Online First: Epub Date]|.

43. Markgraf DF, Klemm RW, Junker M, Hannibal-Bach HK, Ejsing CS, Rapoport TA. An ER protein functionally couples neutral lipid metabolism on lipid droplets to membrane lipid synthesis in the ER. Cell Rep 2014;6(1):44-55 doi:10.1016/j.celrep.2013.11.046[published Online First: Epub Date]|.

44. Benador IY, Veliova M, Mahdaviani K, et al. Mitochondria Bound to Lipid Droplets Have Unique Bioenergetics, Composition, and Dynamics that Support Lipid Droplet Expansion. Cell Metab 2018;27(4):869-85 e6 doi:10.1016/j.cmet.2018.03.003[published Online First: Epub Date]|.

45. Chatzigeorgiou A, Chung KJ, Garcia-Martin R, et al. Dual role of B7 costimulation in obesity-related nonalcoholic steatohepatitis and metabolic dysregulation. Hepatology 2014;60(4):1196-210 doi:10.1002/hep.27233[published Online First: Epub Date]|.

46. Liebisch G, Binder M, Schifferer R, Langmann T, Schulz B, Schmitz G. High throughput quantification of cholesterol and cholesteryl ester by electrospray ionization tandem mass spectrometry (ESI-MS/MS). Biochim Biophys Acta 2006;1761(1):121-8 doi:10.1016/j.bbalip.2005.12.007[published Online First: Epub Date]|.

47. Herzog R, Schwudke D, Schuhmann K, et al. A novel informatics concept for high-throughput shotgun lipidomics based on the molecular fragmentation query language. Genome Biol 2011;12(1):R8 doi:10.1186/gb-2011-12-1-r8[published Online First: Epub Date]|.

48. Herzog R, Schuhmann K, Schwudke D, et al. LipidXplorer: a software for consensual cross-platform lipidomics. PLoS One 2012;7(1):e29851 doi:10.1371/journal.pone.0029851[published Online First: Epub Date]|.

49. Wickham H. Reshaping Data with the reshape Package. J Stat Softw 2007;21(12):1-20

50. Wickham H. ggplot2: Elegant Graphics for Data Analysis. Springer-Verlag New York 2009

